# LifeSciBench: Evaluating Language Models on Realistic, Expert-Level Tasks in the Life Sciences

**DOI:** 10.64898/2026.08.13.744657

**Authors:** Amelia Liu, Andrew Ho, Anne Marie Droste, David Martin, Edmund Wong, Edward Zhou, Isabelle Zhou, Joshua Park, Joy Jiao, Katie-Rose Skelly, Kenny Kim, Jeremy Li, Kevin Rao, Masatoshi Uehara, Max Marion, Nicole Fitzgerald, Rachel Dias, Suyash Shringarpure, Yuan Yuan, Yunyun Wang

## Abstract

We introduce LifeSciBench, a benchmark of 750 expert-authored tasks designed to evaluate whether language models can handle realistic life science research work. The majority of existing life sciences benchmarks have a narrow scope or are purely knowledge-based, and therefore fail to capture the complexity of real-world research, which often involves ambiguities and requires the accurate execution of multiple dependent judgment calls. Additionally, almost all existing benchmarks span at best a small collection of subdomains within the life sciences; there is at present no existing life sciences benchmark with both the requisite breadth and depth required to convincingly measure proficiency in real-world professional research settings. LifeSciBench addresses this gap by spanning seven representative scientific workflows and seven life science domains, with each constituent task paired with a human expert-written rubric. Across five frontier and domain-specialized models, GPT-Rosalind performs best, with a task-weighted mean normalized rubric score of 0.576 and a task-weighted response pass rate of 36.1% (response-level values are first averaged within each task, and the resulting task-level values are then averaged with equal weight). LifeSciBench remains unsaturated, with 171 tasks (22.8%) having no observed passing response from any evaluated model and 261 tasks (34.8%) having a best-model pass rate below 20%. LifeSciBench therefore serves as a high-resolution evaluation of practical scientific reasoning and operational decision-making in the life sciences.

## 1 Introduction

Recent advances in large language models (LLMs) have produced agentic systems that can reason, use software tools, and solve highly specialized domain-specific problems with increasing sophistication. These capabilities are especially relevant to the life sciences, where research progress relies upon a combination of reasoning from ambiguous yet complex evidence, careful experimental design, and precise decision-making in uncertain settings. To be useful to professionals or academics in the life sciences, models must be capable of handling tasks that reflect the structure and constraints of real scientific work. Existing life-science evaluations mostly emphasize factual knowledge retrieval, as with GeneTuring, or concentrate on computational biology and ‘omics analyses, as in BixBench, SpatialBench, scBench, and CompBioBench (Shang et al., 2025; Mitchener et al., 2025; Workman et al., 2025, 2026; Nair et al., 2026). These evaluations are valuable for measuring more bounded analyses, but their narrower task structures provide limited insight into whether models can support the broader range of reasoning-based capabilities and judgment calls required across realistic life-science workflows. Figure 1 summarizes how LifeSciBench constructs and evaluates tasks aimed at this broader setting.

**Figure 1:**
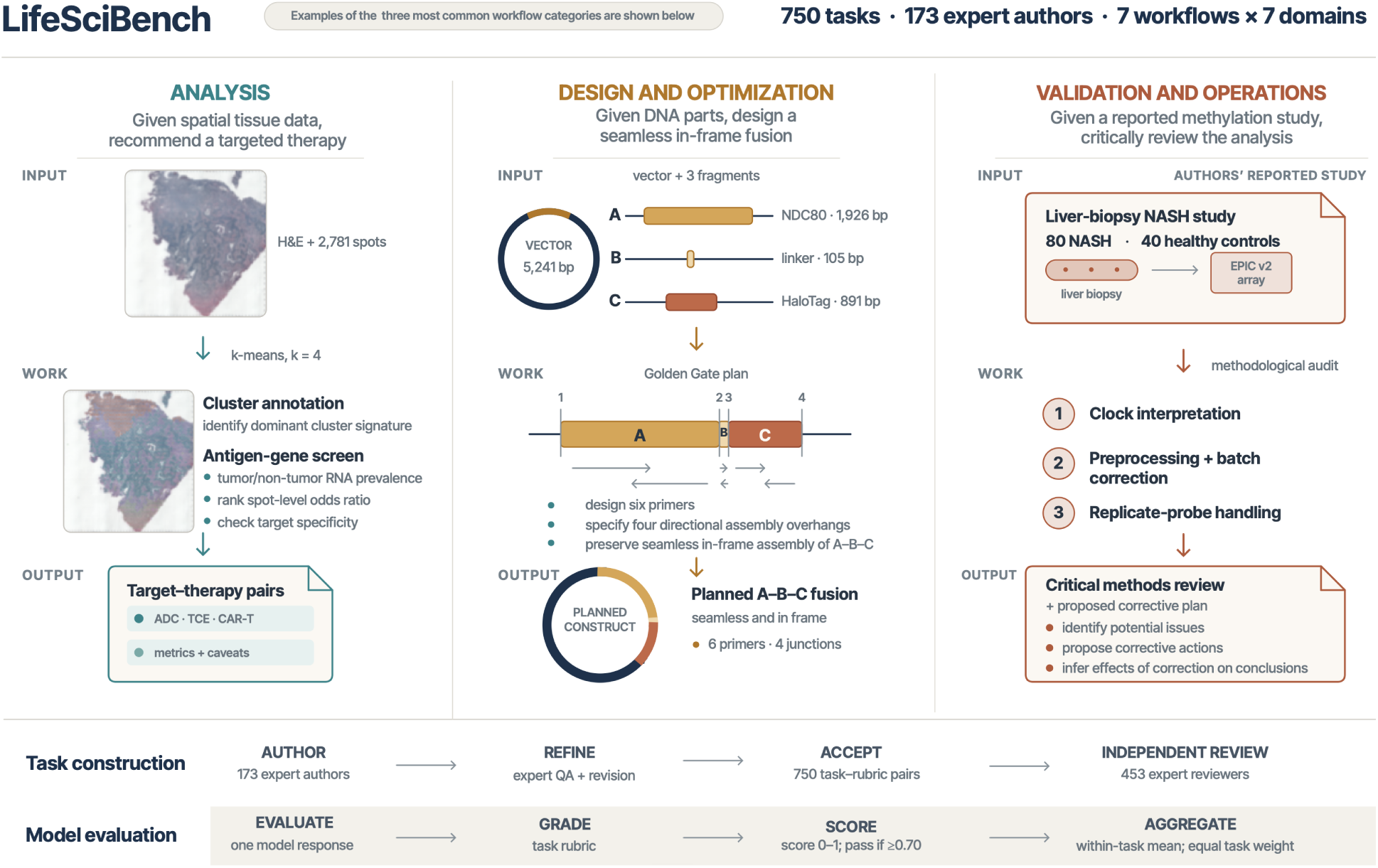
LifeSciBench task examples, construction, and evaluation. LifeSciBench contains 750 task–rubric pairs authored by 173 experts across seven workflows and seven biological domains. The upper panels illustrate three common workflow categories as input–work–output sequences: analysis of spatial tissue data to identify target–therapy pairs; design of a seamless, in-frame NDC80–linker–HaloTag construct and its Golden Gate assembly plan; and methodological review of an EPIC v2 liver-biopsy methylation study, including clock interpretation, preprocessing and batch correction, and replicate-probe handling. The lower panels summarize the benchmark pipeline. Tasks undergo expert quality assurance and revision followed by independent review by 453 experts. For model evaluation, each response is graded against its task-specific rubric, assigned a normalized score from 0 to 1 and a pass indicator at ≥ 0.70, and aggregated using the within-task mean with equal weight assigned to each task.

Accordingly, we present LifeSciBench, an expert-designed benchmark containing 750 expert-level problems designed to evaluate the ability of models to perform frontier-level work in the life sciences. The benchmark is designed around tasks that require a combination of domain-specific reasoning, data analysis, and literature search. Additionally, the benchmark does not only evaluate the ability of the model to produce the correct analysis in response to a well-specified question; rather, the questions are intentionally left somewhat open-ended, such that LifeSciBench also measures the model’s understanding of the appropriate level of detail to supply in its responses.

LifeSciBench is designed to capture aspects of real-world research work that are often missing from existing evaluations:

1. **Reasoning over complex artifacts:** LifeSciBench requires models to reason over artifacts commonly encountered in research settings, such as images, documents, sequence files, molecular structures, and web references.
2. **Situational ambiguity:** In practice, most user interactions with LLMs are not written in an unambiguous and precise textbook-like manner; indeed, the typical interaction more closely resembles human-collaborator interaction, where ambiguities may be present. As such, LifeSciBench aims in part to evaluate whether models can correctly interpret ambiguous context and make evidence-based recommendations from incomplete or conflicting evidence.
3. **Realism:** Tasks in LifeSciBench are organized around the actual functions that scientists perform in practice, from hypothesis generation and data generation to evaluating translational risk and communicating findings to others.
4. **Expert-authored and independently reviewed:** Each LifeSciBench task and accompanying rubric was written by a domain specialist and reviewed through a multi-round quality assurance (QA) process.

### 1.1. Related Work

Recent life science benchmarks have expanded beyond factual biology question answering and now tend to differ in the particular type of research capability they assess. One line of work evaluates research-assistant behaviors such as literature retrieval, database lookup, figure and table interpretation, and reasoning about bench protocols. For instance, LAB-Bench introduced a broad suite of biology research tasks along these lines (Laurent et al., 2024); LABBench2 extended this with more realistic, open-response settings, including tasks involving patents, clinical trials, and “messy” data (Laurent et al., 2026). These benchmarks aim to evaluate performance on practical research tasks, but they still fail to thoroughly evaluate expert scientific judgment — indeed, in practice, nearly all research decisions require not only the identification or manipulation of biological information, but also the judgment-heavy process of weighing imperfect or incomplete evidence, reasoning within experimental constraints, and the production of clear, actionable outputs.

A second line of work evaluates agentic capabilities in computational biology. For example, BixBench tests LLM agents on bioinformatics scenarios requiring code execution, dataset exploration, and multi-step analysis, but is mostly saturated by current frontier models (Mitchener et al., 2025). More recently, GeneBench and its expanded successor, GeneBench-Pro, evaluate models on unusually challenging, multistage quantitative biology analyses involving noisy data, quality control, statistical modeling, confounding, and downstream inference (Li and Ho, 2026; Li et al., 2026). Their use of synthetic data and extensive ablations is intended to make the graded target identifiable and to distinguish defensible analyses from plausible but incorrect workflows. These benchmarks are valuable in that they move beyond static question answering and require models to reason over data of unknown provenance, but their scope is primarily concentrated in computational biology.

LifeSciBench is closer in spirit to the former line of work, and addresses the remaining gap by requiring a combination of expert-level scientific reasoning and data analysis across a broad swathe of applied life science research. Its constituent tasks span multiple subfields of biology and stages of the drug discovery process. Additionally, rather than evaluating only binary correctness, LifeSciBench uses expert-authored rubrics to assess whether models reach conclusions through scientifically valid reasoning, appropriate consideration of relevant evidence, and operationally useful decision-making. Thus, LifeSciBench allows models to be scored both on their intermediate reasoning abilities as well as on their final conclusions.

## 2 Benchmark Construction

### 2.1. Benchmark Taxonomy & Coverage

Life scientists perform a wide range of tasks in day-to-day work, ranging from highly specialized, domain-specific work to more routine activities shared across multiple domains. We began by surveying practicing scientists about the workflows they most often utilize in applied research settings. We then grouped their responses into seven categories which describe approximately distinct types of workflows (Table 1). We also categorized each task into seven biological domains and seven data-source and evidence categories (Table 1). Note that while we use these dimensions to perform coverage and subgroup analyses, they are high-level descriptors rather than a strict decomposition of abilities: realistic tasks may combine multiple capabilities and evidence types.

**Table 1:**
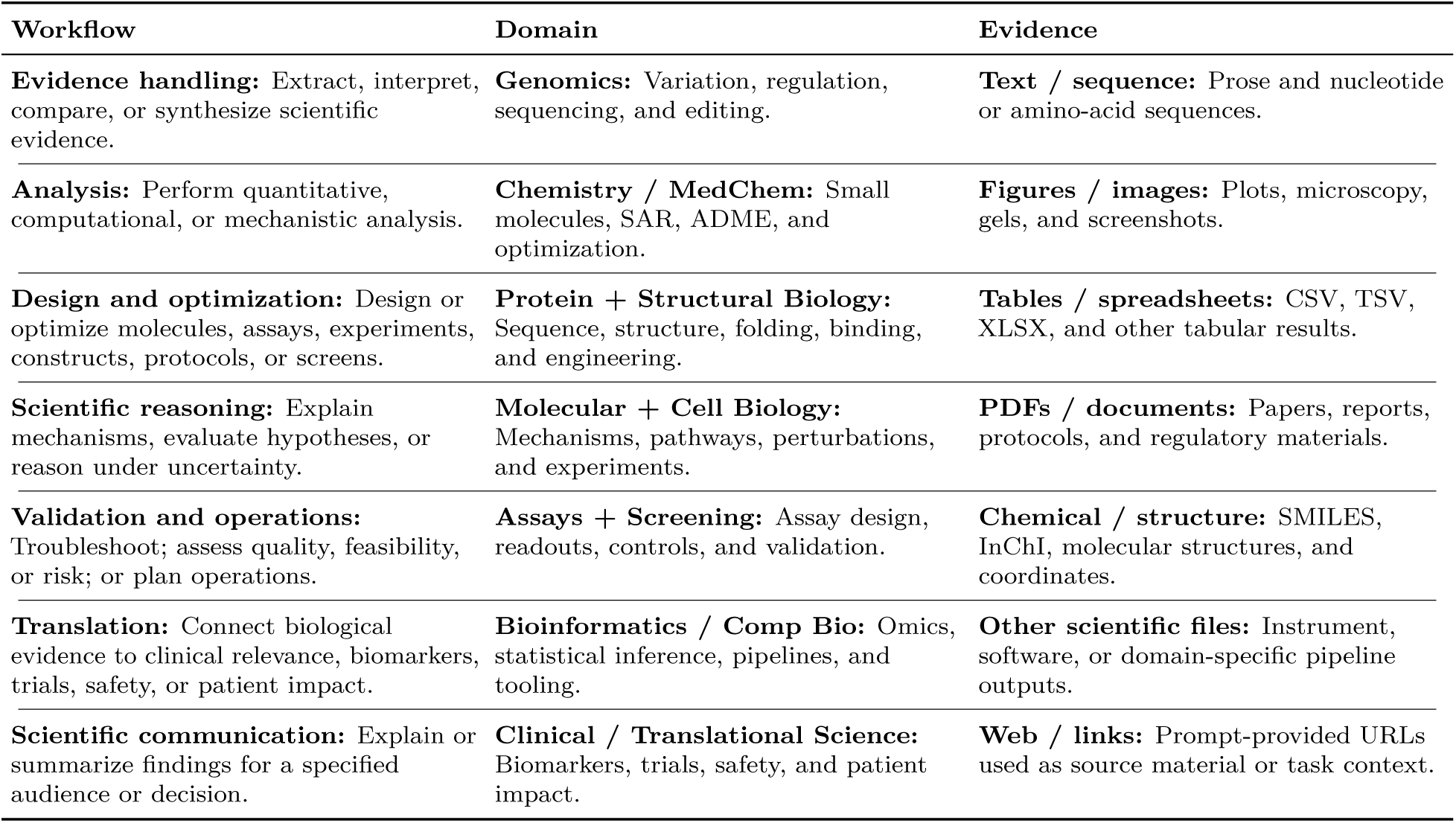
Complementary task taxonomies in LifeSciBench. Workflows describe the primary research activity, domains the scientific setting, and evidence types the source material provided or referenced. The columns are independent: horizontal alignment does not imply a relationship between entries in the same row.

The problems in LifeSciBench span both computational and experimental contexts, reflecting the need for models to move between different forms of evidence and modes of analysis. Figure 2 shows their joint distribution.

**Figure 2:**
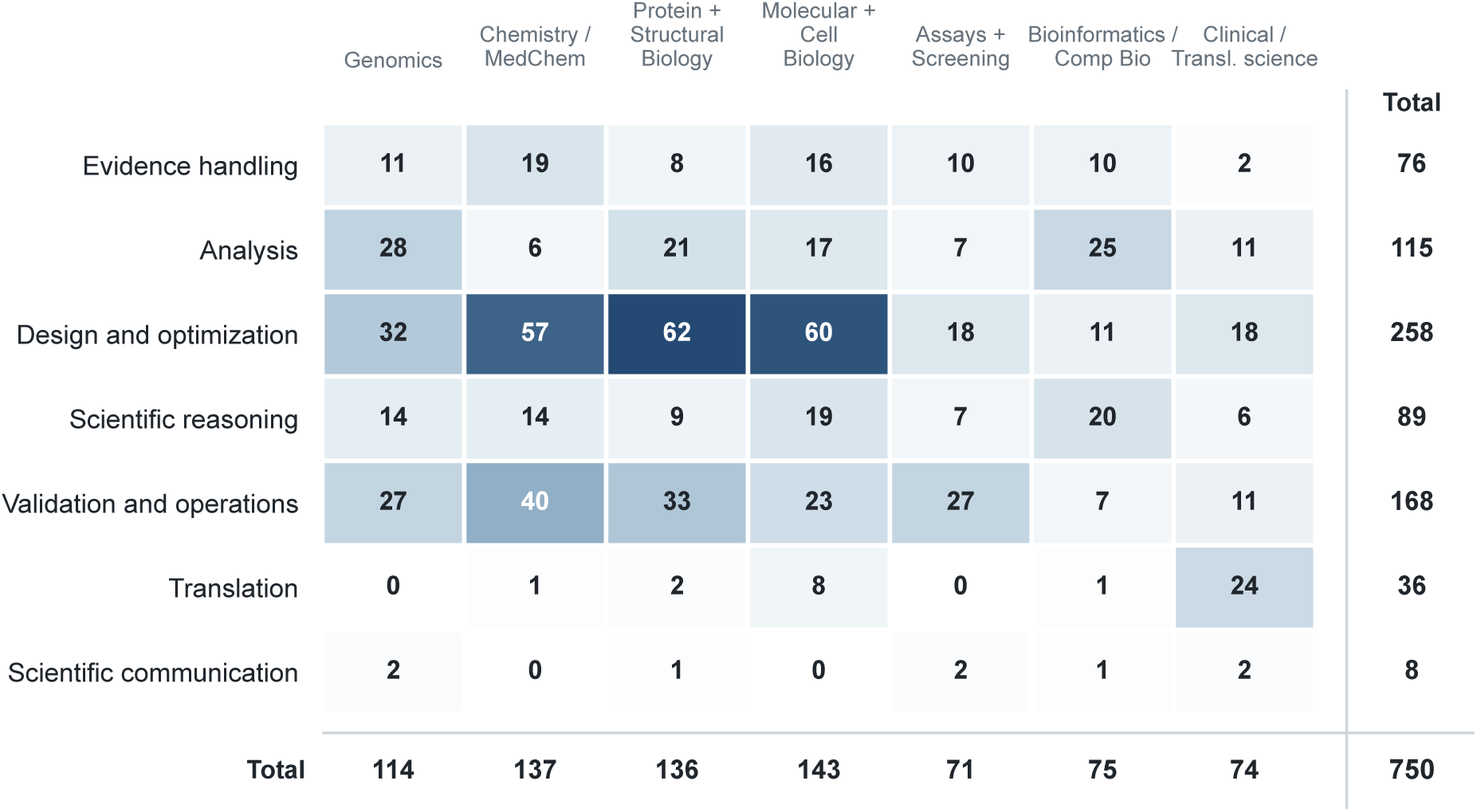
LifeSciBench coverage across workflows and biological domains. Cells show task counts for each workflow–domain combination, for a total of 750 tasks. Workflows describe the primary research activity the task is supposed to reflect, whereas domains denote the scientific setting. Individual tasks may require capabilities spanning multiple categories.

### 2.2. Expert Writer Cohort

The tasks in LifeSciBench were created by 173 expert scientists spanning a diverse range of life science disciplines chosen to ensure that the benchmark reflects the full breadth of expertise needed to evaluate agentic AI systems across life science research. Experts were required to have completed a Ph.D. in a relevant discipline, such as biochemistry, molecular biology, neuroscience, immunology, pharmacology, medicinal chemistry, computational biology, or a related field, and have at least two years of experience as practicing scientists in the biotechnology or pharmaceutical industries. To ensure that tasks more closely resembled real-world work rather than textbook questions, we selected contributors whose collective experience spanned computational, experimental, translational, and clinical research.

### 2.3. Task Structure

Each LifeSciBench task consists of a prompt containing a scientific question, any supporting artifacts (*e.g.*, images, assay results, sequence data), and a grading rubric. Below, we describe the nature of each of these components.

#### Questions

Each question in LifeSciBench comprises a prompt written in natural scientific language, structured as a scientist might pose a problem to a knowledgeable colleague or assistant, and asks a scientific question which the model must answer in free text. The questions range from focused, single-answer queries such as identifying the germline of a humanized antibody or computing a combinatorial count, to multi-step analytical tasks requiring explicit reasoning and scientific judgment.

**Figure 3:**
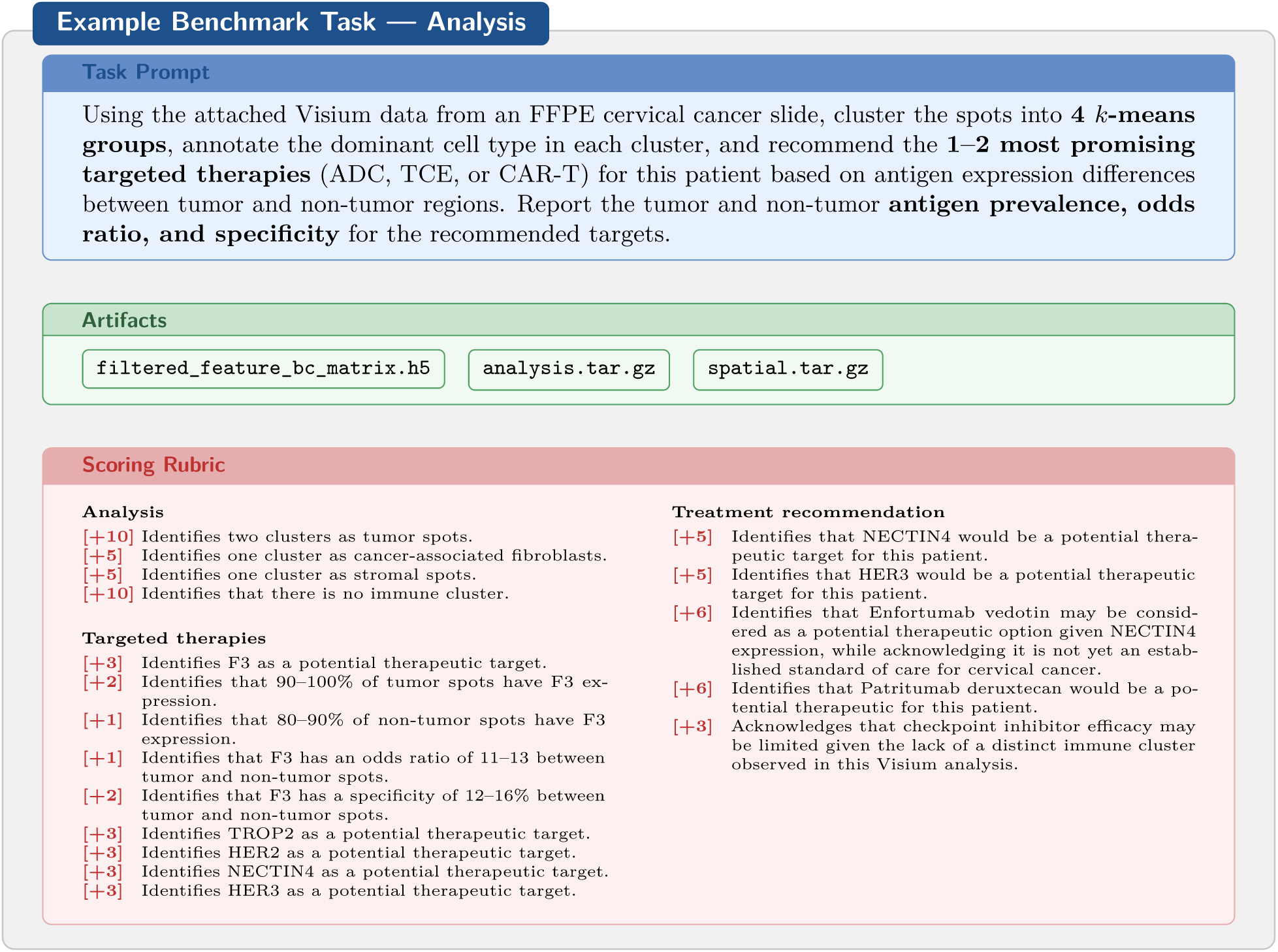
Example LifeSciBench task. Each task comprises an expert-written prompt, a set of artifacts or contextual evidence, and a fine-grained evaluation rubric. A representative subset of rubric criteria is shown.

The tasks cover core research capabilities such as evidence interpretation, data analysis, experimental design, troubleshooting results, and scientific communication. Tasks also test practical behaviors, such as complex instruction-following, incorporating heterogeneous context, and uncertainty handling.

#### Artifacts

Many questions require analysis of task-specific artifacts, which include molecular representations such as SMILES or InChI strings, nucleotide or amino-acid sequences, tabular datasets, PDFs, raw instrument outputs, microscopy images, gel images, experimental figures, or other scientific files.

#### Evaluation

Evaluation is conducted in a single-turn setting. Each model receives the prompt and any associated artifacts once and produces one final response to the original question. Although multi-turn settings are arguably more reflective of real usage, a single-turn design, which is currently the standard in benchmark construction, isolates task-level performance on self-contained scientific problems while preserving realistic complexity in the prompt, supporting context, and expected response. All models had access to the same Internet-browsing-enabled, containerized scientific computing environment (Section 4.1).

#### Rubrics

LifeSciBench uses task-specific rubrics to evaluate model responses. For each task, rubric criteria describe attributes of a response that should be rewarded or penalized. Rubric criteria range from specific facts that should be mentioned in the response, to explicit reasoning steps the model must demonstrate, to quantitative outputs evaluated within an accepted tolerance.

To support criterion-level analyses, each rubric criterion was also assigned two descriptive labels: an *operation* label, describing what the response must do, and an *answer-form* label, describing the nature of the response. These labels are distinct from the task-level workflow, domain, and evidence taxonomies because a single task may contain criteria spanning multiple operations and answer forms. The labels were produced by an automated rubric-item classifier and were used only for descriptive subgroup analyses. Appendix Table 3 lists the complete taxonomy.

In order to calculate partial credit, for each response, signed rubric contributions are divided by the task’s total positive points and clipped to [0, 1]; see Section 4.2 for the precise definition. We chose to score tasks in this manner because we wished to capture the richness of models’ intermediate reasoning that would otherwise be lost if we were to solely compare a final answer to a single deterministic value: indeed, in scientific work, an answer may depend on whether the model uses the correct evidence, states relevant assumptions, applies appropriate methods, respects task constraints, and communicates conclusions at the right level of certainty.

This criterion-level scoring framework allows us to measure both final task success and the component capabilities that contribute to that success. Table 2 summarizes benchmark scale, supporting evidence, task complexity, and rubric granularity.

**Table 2:** LifeSciBench scale and task composition.

| Benchmark Characteristic | Value |
| --- | --- |
| Total tasks | 750 |
| Task attachments | 1,009 |
| Tasks with one or more attachments | 395 (52.7%) |
| Tasks with prompt-provided URLs | 37 |
| Tasks requiring multiple reasoning or decision-making steps | 79% |
| Average reasoning / decision-making steps per task | 4 |
| Expert-written rubric criteria | 19,389 total (25.9 per task) |

### 2.4. Review Process

All tasks underwent a multi-stage expert review process. Tasks could undergo as many revision cycles as needed before acceptance. Accepted tasks averaged six self-directed automated review cycles and completed at least two rounds of expert review.

In the process of expert review, each task was explicitly checked against four criteria:

- **Task-rubric consistency:** Reviewers ensured consistency between the question and rubric. In particular, they checked that rubric criteria were actually requested by the question, that the rubric did not introduce unasked-for requirements, and that criteria could be evaluated objectively rather than being dependent on the specific task author.
- **Realism:** Reviewers ensured that tasks reflected real-world scientific work and explicitly tested multi-step reasoning beyond memorization or simple lookups. Reviewers checked that the task required meaningful scientific judgment, evaluation of evidence, data analysis, or design understanding.
- **Scientific fidelity:** Reviewers ensured that the scientific conclusions and assumptions were verified to be accurate. Rubric items were required to be supported by provided evidence, accepted scientific consensus, or expert judgment. Reviewers also checked that tasks did not overgeneralize, overstate certainty, or introduce unsupported assumptions.
- **Spelling, grammar, and formatting:** Questions and rubrics were reviewed to remove spelling, grammar, and formatting errors that could interfere with task interpretation or grading.

**Figure 4:**
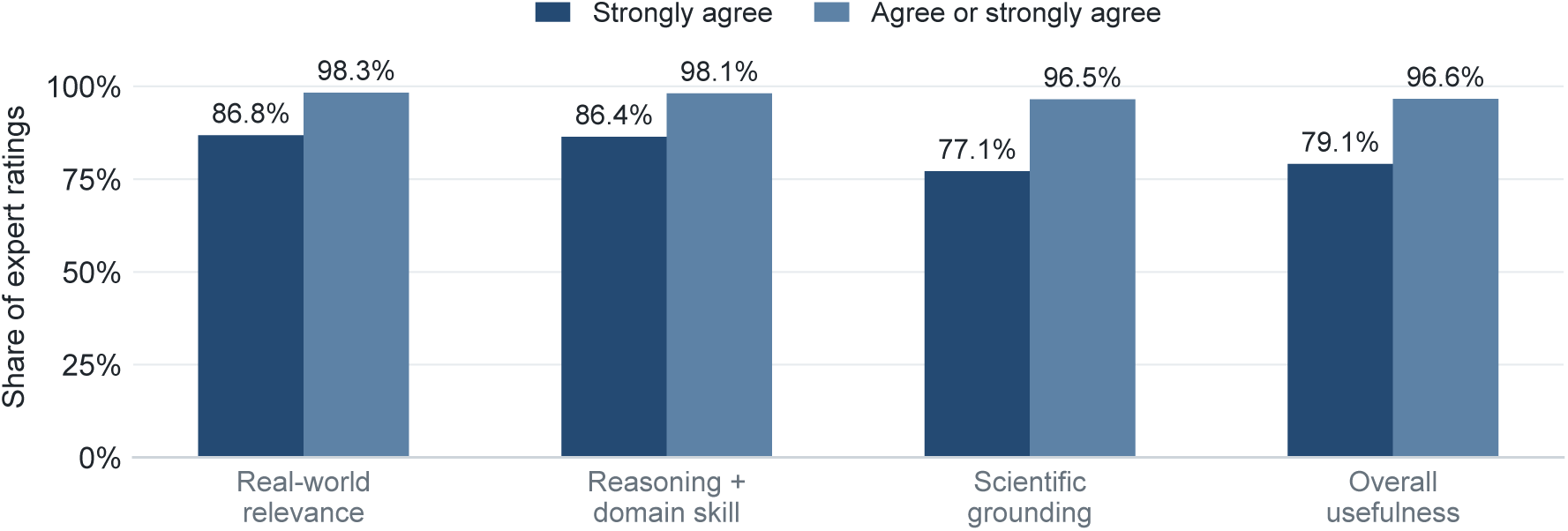
Independent expert validation results. Bars show the reported share of expert ratings that selected “strongly agree” and the cumulative share that selected “agree” or “strongly agree” for each validation criterion.

The criteria for a task’s inclusion in LifeSciBench required either a verifiable answer or expert consensus, the latter corresponding to at least 90% agreement among domain experts. This process was designed to ensure that LifeSciBench tasks are scientifically rigorous, consistently gradeable, and representative of realistic life science work.

## 3 Benchmark Validation

To validate the scientific quality and utility of LifeSciBench, we conducted an independent expert review of the benchmark tasks. This validation was separate from the task construction and review process described above. Reviewers were distinct from the task writers and had substantial life science expertise: a total of 453 expert reviewers participated, of which 97% held a Ph.D. or equivalent; on average, reviewers had 12 years of field experience and 14 peer-reviewed publications, and 88% reported receiving at least one award or fellowship.

Reviewers were asked to assess whether each task reflected the following four high-level properties (1) real-world relevance, (2) scientific reasoning and domain-skill alignment, (3) scientific grounding, and (4) overall usefulness for assessing model performance.

Across reviewed tasks, expert ratings were high on all four dimensions. For real-world relevance, 86.8% of ratings were (on a Likert scale) “strongly agree” and 98.3% were “agree” or “strongly agree” that tasks reflected realistic life science work. For scientific reasoning and domain-skill alignment, the corresponding percentages were 86.4% and 98.1%; for scientific grounding, 77.1% and 96.5%; and for overall usefulness, 79.1% and 96.6%.

Overall, these results support the conclusion that practicing experts consider LifeSciBench tasks realistic assessments of the reasoning and scientific judgment required in applied life science research.

## 4 Experimental Setup & Scoring

We evaluated a set of frontier general-purpose and domain-specialized language models on LifeSciBench: GPT-5.4, GPT-5.5, GPT-Rosalind, Gemini 3.1 Pro, and Grok 4.3. All models were evaluated in a single-turn setting: each model receives the task prompt and any associated artifacts and must produce a final answer without follow-up interaction.

### 4.1. Evaluation Protocol

For each task, we provided the model under evaluation with the prompt and associated context. For tasks involving artifacts, the model was given access to the relevant files. Models were instructed to answer the task directly and to include its reasoning and key calculations alongside any caveats or assumptions made for the final answer. Internet access was enabled and unrestricted in the evaluation environment.

All model runs used the same Linux-based scientific computing Docker image. The container provided standard scientific-computing libraries including NumPy, SciPy 1.14.1, pandas, scikit-learn 1.8.0, statsmodels 0.14.6, Matplotlib 3.10.8, h5py 3.15.1, and openpyxl 3.1.5; bioinformatics and omics tools including BLAST+, bedtools, BWA, Bowtie 2, MAFFT, MMseqs2, MUSCLE, PLINK2, Scanpy 1.11.5, Seurat, DESeq2, edgeR, limma, pysam 0.23.3, and pybedtools 0.12.0; and chemistry, structural-biology, and simulation tools including RDKit, Open Babel, OpenMM 8.4.0.post2 on AMD64 or 8.5.0b0 on ARM64, MDTraj 1.11.0, Gemmi 0.7.3, DSSP 4.5.6, PyMOL, APBS, CP2K, GROMACS, LAMMPS, NWChem, and Quantum ESPRESSO. Bioconductor was fixed at version 3.22. Packages without a listed version were included without a version pin.

Model outputs were automatically graded against the expert-written rubric associated with each task using GPT-5.5 as a LLM judge. For each response, the judge received the task question text, expert-written rubric, and model’s final answer, but not attachments or tool transcripts. It assigned a value between zero and one to each rubric criterion according to the task-specific scoring scheme. We aggregate these values into a normalized rubric score between 0 and 1, then summarize response scores and pass rates within tasks before computing overall model aggregates (tasks were equally weighted).

### 4.2. Metrics

We report model performance on a given task using two complementary metrics: the normalized rubric score and task pass rate. Both begin with the normalized response-level score, which can be considered the amount of partial credit a response receives (from 0 to 1). A response is considered passing when its normalized rubric score is *≥* 0.70.

#### Criterion Satisfaction

For each rubric criterion in a response, the grader assigns a satisfaction value *s_qrj_ ∈* [0, 1], where *q* indexes tasks, *r* indexes sampled responses for task *q*, and *j* indexes rubric criteria within that task. A value of 0 indicates that the response does not satisfy the criterion, 1 indicates full satisfaction, and intermediate values indicate partial satisfaction. For a positive criterion, *s_qrj_* is the fraction of the criterion’s points awarded; for a negative-penalty criterion, it represents the extent to which the penalized behavior is present. Thus, for example, a satisfaction value of 0.8 on a five-point positive criterion contributes four points to the response’s rubric score.

#### Normalized Rubric Score

We calculate the normalized rubric score for each model response directly from the criterion satisfaction values defined above. This score is the fraction of the task’s available positive points earned by a response after any penalties are applied. Each rubric item has a positive or negative point value, and the grader assigns it a value between 0 and 1 based on how closely the response matches that item. We multiply each item’s value by its point value, add the results across all items, and divide by the total positive points available for the task. Positive-point items add credit when they are met, whereas negative-point items subtract credit when the penalized behavior is present. We then limit the final score to the range from 0 to 1. For task *q* and response *r*, this is

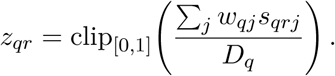

Here, *w_qj_*is the point value of rubric item *j*, *s_qrj_* is its 0-to-1 grader value, and *D_q_*is the total positive points available for the task. Intuitively, this can just be considered the degree to which the response satisfies the rubric, where 0 is when the response satisfies none of the criteria and 1 is when the response satisfies all of the criteria.

#### Task Pass Rate

A sampled response is considered passing when its normalized rubric score *z_qr_* is *≥* 0.70. For each model/task pair, we first average the response scores and pass indicators over all runs with that model/task combination. We then average those task-level quantities across observed tasks with every task weighted equally. Thus, the reported task pass rate is the mean within-task response pass fraction.

For the task-level subgroup analyses by workflow and artifact setting later described in this manuscript, we apply the same normalized rubric score and pass rate definitions. For the criterion-level analyses later described, we converted all criteria to a scale on which higher values indicate better performance; namely, for criteria that award points, we used the grader’s satisfaction value directly. For criteria that deduct points, we used one minus the satisfaction value, such that avoidance of the penalized behavior results in a higher score. For each model and task, we then computed a weighted average of the criteria within each operation or answer-form category where each criterion was weighted by the absolute number of rubric points assigned to it. Finally, we averaged the resulting category scores across tasks.

## 5 Results

Across LifeSciBench, performance varies substantially by task type, workflow, and response format. Overall, GPT-Rosalind performed best, with a mean normalized rubric score/pass rate of (95% task-bootstrap intervals in parentheses) 0.576 (0.560–0.591) / 36.1% (33.5–38.8%). GPT-5.5 scored 0.519 (0.505–0.534) / 25.7% (23.4–28.0%); Gemini 3.1 Pro, 0.515 (0.499–0.531) / 23.6% (21.0–26.2%); GPT-5.4, 0.479 (0.464–0.493) / 20.7% (18.6–22.8%); and Grok 4.3, 0.399 (0.383–0.416) / 13.0% (11.0–15.1%) (Figure 5A–B).^1^ We obtained these intervals by resampling tasks with replacement 10,000 times, recomputing the mean score and pass rate in each replicate, and taking the 2.5th and 97.5th percentiles. GPT-Rosalind also displays the highest per-task mean score on 386 of 750 tasks.

Pass rates remain modest across all models, indicating that even frontier models do not consistently complete tasks in a satisfactory manner. We quantify the headroom to saturation for LifeSciBench by computing the task pass rate achieved by any evaluated model on each task: Figure 5C groups these values into mutually exclusive bins, showing where tasks fall from no passing sampled responses to high best-model pass rates.

**Figure 5:**
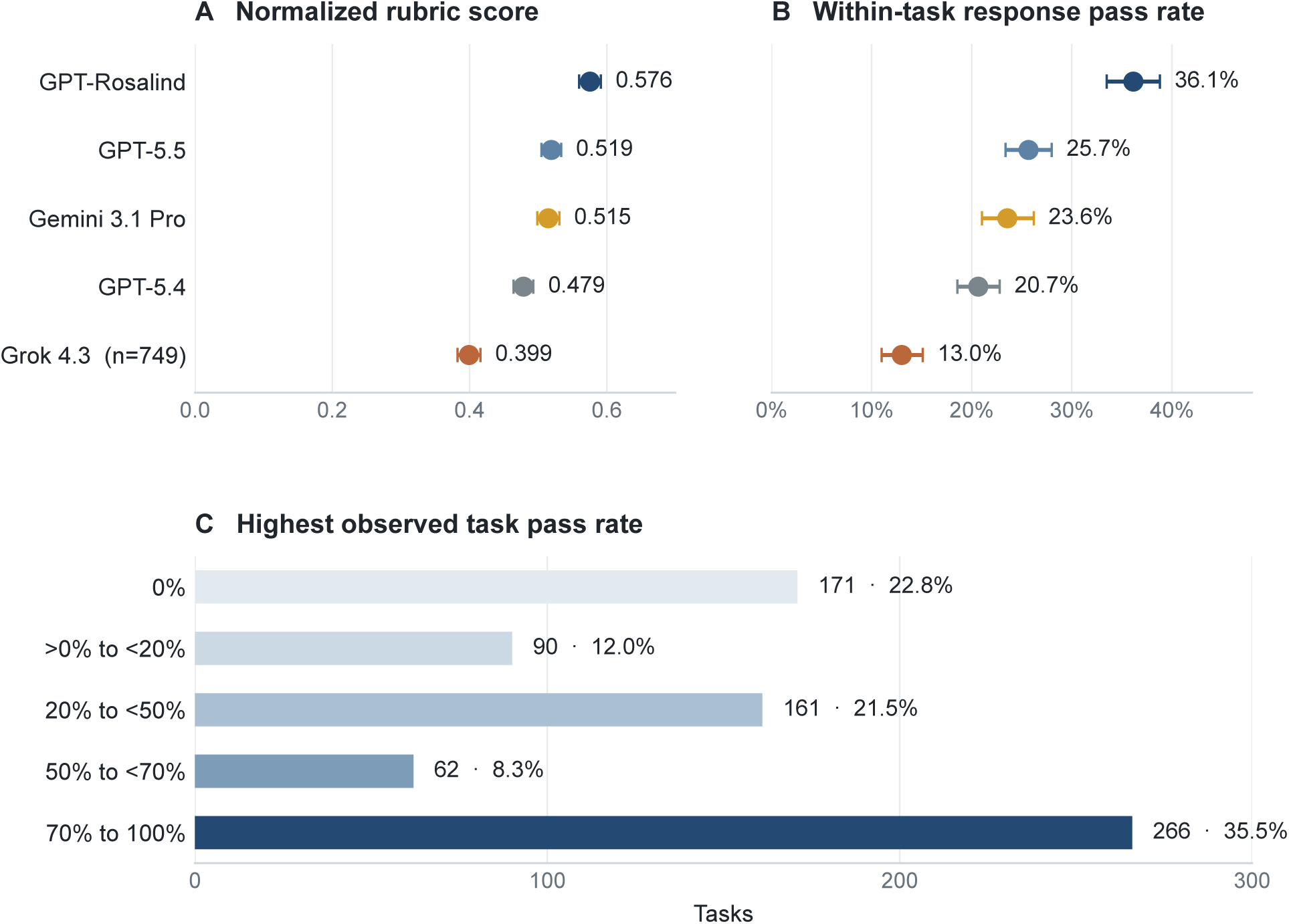
Overall LifeSciBench performance and benchmark headroom. **A–B,** Points show the task-weighted mean normalized rubric score and mean within-task response pass rate for each model; intervals are 95% percentile intervals from 10,000 bootstrap resamples of tasks. C, Distribution across tasks of the highest valid-response pass fraction observed among the evaluated models. Bins are mutually exclusive; labels give the task count and share of the overall benchmark.

Under this binning scheme, 422 tasks (56.3%) have a best-model pass rate below 50%, including 261 tasks (34.8%) with a best-model pass rate below 20%, indicating that LifeSciBench remains far from saturated and retains headroom for measuring future model progress. The lowest, below-20% bin is concentrated in Design & Optimization and Analysis, which together account for 60.9% of tasks in that range.

We next examine where the various models perform best within LifeSciBench. Since aggregate scores can obscure substantial variation across different dimensions, we analyze performance both at the model level and across task categories.

### 5.1. Model Strengths

Overall, within-model performance was relatively similar across workflow types (Figure 6), though Translation was the best-performing subgroup among the well-represented workflows (i.e. excluding Scientific Communication); tasks in this workflow category require models to connect preclinical or biological evidence to downstream clinical or translational relevance. For example, GPT-Rosalind achieved a mean score of 0.712 across the 36 Translation tasks, compared with its overall mean score of 0.576.

**Figure 6:**
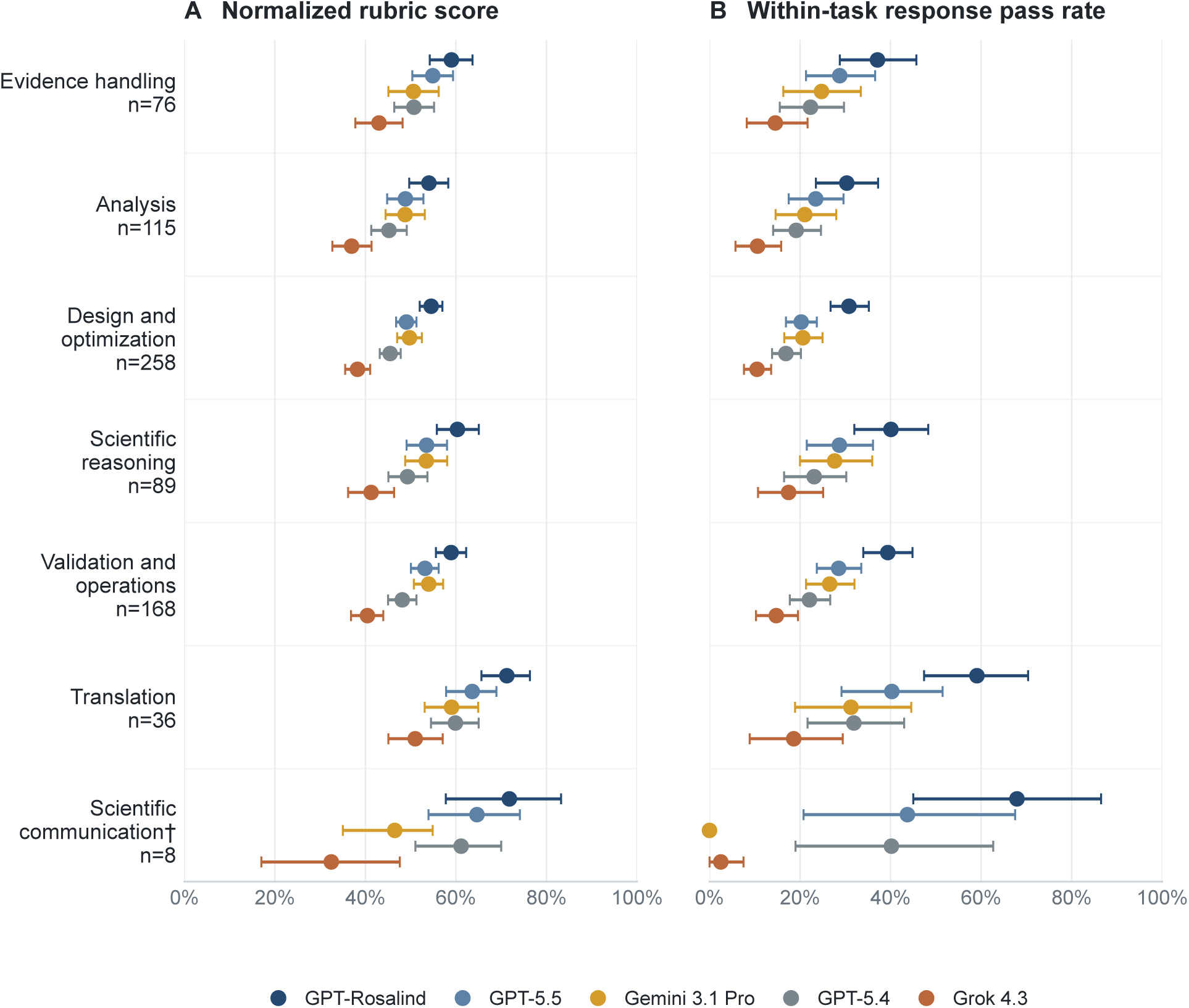
LifeSciBench performance by scientific workflow. Points show task-weighted mean normalized rubric score and mean within-task response pass rate within each workflow. Intervals are 95% percentile intervals obtained by resampling tasks. Workflow labels report full benchmark subgroup sizes; Grok has results for 167 of the 168 Validation and operations tasks.

Rubric-level results suggest that this relative strength extended broadly across components of translational reasoning rather than being driven by a single component. After weighting criteria by rubric points within each task and then weighting tasks equally, GPT-Rosalind’s mean scores for mechanism explanation, critique/validation, and evidence interpretation were 0.721, 0.752, and 0.773 within Translation, compared with 0.602, 0.667, and 0.669 for the same rubric types in other workflows.

#### 5.1.1. Remaining Gaps

On the other hand, performance on tasks involving artifacts was significantly lower across models than on text-only tasks: for example, GPT-Rosalind achieved a 44.6% pass rate on text-only tasks versus 28.6% on tasks with attached artifacts. GPT-5.5 shows the same qualitative pattern, with pass rates of 29.5% and 22.2%, respectively.

**Figure 7:**
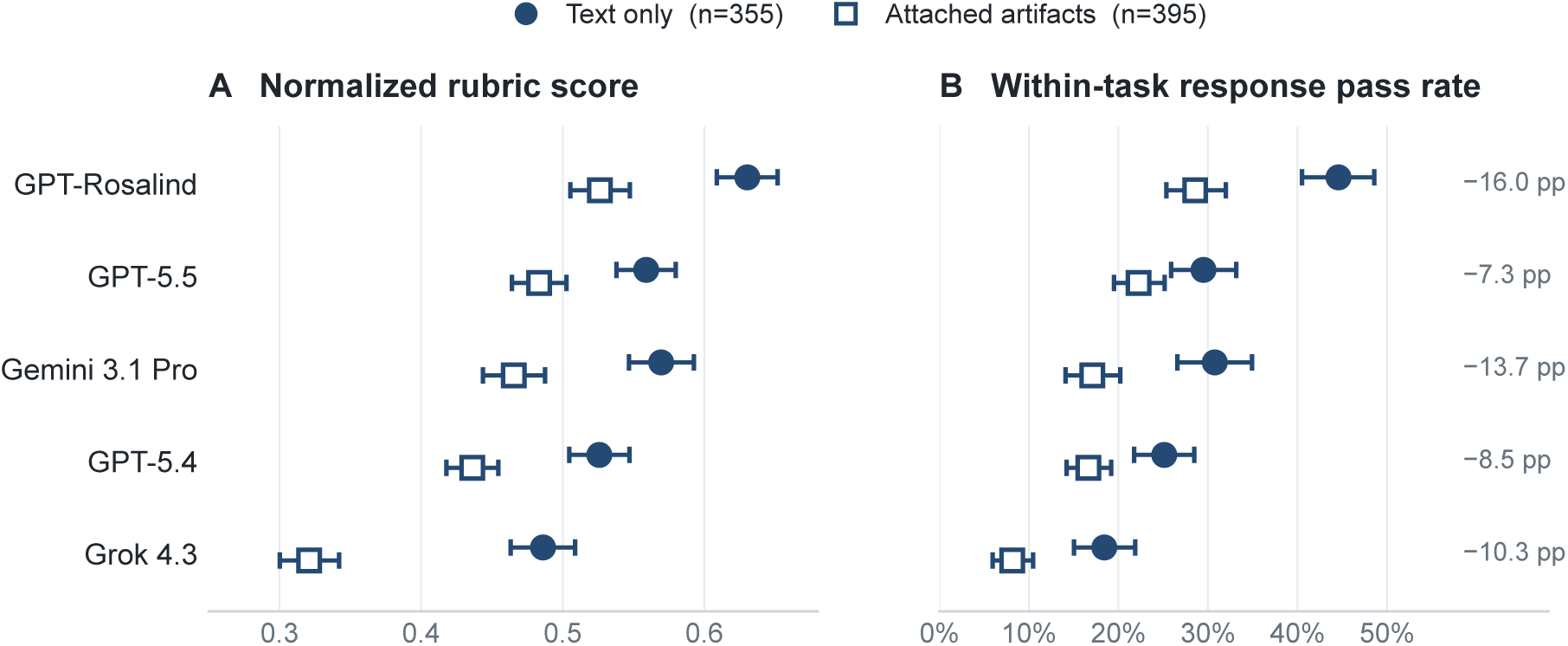
Model performance by evidence setting. Points show a comparison of task-weighted mean scores and mean within-task response pass rates for text-only tasks and tasks that involve artifacts. Intervals are 95% percentile intervals obtained by resampling tasks. Right-side labels give the observed pass-rate difference. Grok has results for 394 of the 395 attached-artifact tasks.

To test whether this gap reflected differences in other characteristics that differed between artifact and text-only tasks, we tested two complementary regression adjustments for each tested model: (1) a task-level adjustment for differences between entire tasks, and (2) a criterion-level adjustment for differences between individual grading requirements. In both analyses, “adjustment” means treating characteristics of the task or criteria as covariates in the regression, such that the performance gap between artifact-bearing and text-only problems is estimated only after accounting for those characteristics.

The task-level regression adjustment included the following covariates: workflow type, each task’s dominant operation, the requested form of the answer, the required exactness of the response, reasoning depth, the number of rubric criteria, and prompt length. This adjustment process reduced the task-level pass-rate gap by 18–38%, but the adjusted gap remained negative for every model (Appendix Figure 10A).

The criterion-level adjustment included the following covariates: workflow, each criterion’s operation, the requested form of the answer, the required exactness of the response, and reasoning depth. This adjustment process changed the raw artifact-associated gaps by at most 1.2 percentage points, leaving adjusted gaps of *−*15.6 to *−*7.0 percentage points (Appendix Figure 10B). These results, while purely observational, indicate that the performance gap persists even after accounting for task and rubric categories.

Criteria requiring models to return precise biological or chemical outputs were also a shared area of lower performance across models. Two related rubric categories capture this requirement: the answer-form category “sequence/structure” (describing the form of the expected answer), and the operation category “generate/construct”. These categories overlap substantially: 841 of 922 generate/construct criteria (91.2%) were also labeled sequence/structure.

Using the weighted criterion score defined above, “sequence/structure” was the lowest-scoring of the nine answer-form categories for every model, ranging from 0.402 for GPT-Rosalind to 0.244 for Grok 4.3 across 145 contributing tasks. “Generate/construct” was likewise the lowest-scoring of 12 operation categories for every model, ranging from 0.418 to 0.219 across 125 tasks. This pattern is practically important, as many life-science workflows require directly usable outputs, such as nucleotide sequences, molecular constructs, or chemical structures.

#### 5.1.2. Models Consistently Achieve Partial Credit Without Full Task Success

Rubric-level scores also reveal substantial partial progress even while the binary pass rates remain low, illustrating the fact that models often are able to identify relevant evidence or complete part of the reasoning path while remaining unable to fully complete a task. For example, for GPT-Rosalind, 109 tasks have pass rates below 20% while still receiving at least 50% rubric score (Appendix Figure 8). This phenomenon demonstrates the utility of reporting both the normalized rubric scores as well as the binary pass rates—while the latter is more indicative of a model’s readiness to directly substitute for human researcher labor, the former allows a more fine-grained view of models’ relative and partial capabilities, which remain useful.

### 5.2. Model Specific Performance Profiles

GPT-Rosalind is the strongest overall model, but task-level results reveal meaningful variation in where models excel. For example, GPT-5.5 and Gemini 3.1 Pro are close in aggregate performance, with mean rubric scores of 0.519 and 0.515 and pass rates of 25.7% and 23.6% respectively.

However, their task-level profiles differ: across all models, Gemini was the leader on 214 tasks despite its slightly lower aggregate score, whereas GPT-5.5 was the leader on only 61 (Appendix Figure 11). Examining the pairwise correlations between different models’ task-level scores revealed this as a consequence of the fact that GPT-family models displayed much more strongly correlated performance with one another than with other models. Specifically, the correlation between task-wise scores between GPT-5.5 and GPT-Rosalind was *r* = 0.93, whereas that between Gemini and GPT-Rosalind was *r* = 0.64.

Thus, GPT-5.5 and GPT-Rosalind tended to perform well or poorly on the same tasks, whereas Gemini exhibited a qualitatively different pattern of strengths and weaknesses, consistent with Gemini leading on a substantial set of tasks even without a higher overall mean score. More generally, these results suggest that model families differ not only in average performance but also in their relative strengths across the categories of scientific capabilities present in LifeSciBench. For example, GPT-5.5 consistently performed better on problems involving experimental design, interpretation of evidence, and scientific critique or validation.

These results illustrate the utility of reporting performance results at various levels of aggregation as well as at both the rubric score and binary pass rate level.

## 6 Discussion

In this manuscript, we introduced LifeSciBench, a benchmark of 750 expert-authored problems in the life sciences spanning seven types of scientific workflows and seven biological domains. LifeSciBench measures whether models can produce scientifically grounded, operationally useful responses to realistic research tasks.

Each task within LifeSciBench was developed using a multi-round process of task development and refinement, review, and validation involving practicing life scientists with both Ph.D.-level training and industry experience across a variety of domains. Tasks are graded using detailed expert-written rubrics totaling 19,389 criteria, designed to evaluate not only final correctness but also the quality of intermediate reasoning and communication by the model.

Results from evaluating five model systems on LifeSciBench reveal that while current frontier models consistently achieve partial solutions, they remain unable to consistently complete realistic research tasks end-to-end, with the highest-scoring model, GPT-Rosalind, achieving an average task-level pass rate of 36.1% over all problems. Results stratified by model and task categories further reveal both performance trends common across all models as well as differential strengths between model families. For example, all models tended to display relatively strong performance on Translational tasks relative to other task categories, while the GPT family of models consistently performed better than Gemini on problems with components of experimental design and evidence interpretation.

With respect to limitations, LifeSciBench is designed to measure model performance on realistic, self-contained life-science tasks, but it is ultimately still a constructed evaluation without the complexities of real-world research problems. While we focus on core workflows that are broadly relevant across life-science research, due to the tremendous complexity and breadth of the life sciences, it is not feasible for a single benchmark to cover every single type of problem that scientists might practically encounter, or to capture the full range of ambiguities present in actual research scenarios, in which there may often not be a single “correct” answer.

Furthermore, the evaluation was conducted in a single-turn setting, where each model receives a task and any associated artifacts then produces one final response. In contrast, real usage of LLMs is almost always multi-turn and involves user interaction such as requests for clarification of uncertain points or user-driven course corrections.

Accordingly, performance on LifeSciBench should be interpreted as evidence of one-shot task-level capability under realistic scientific constraints rather than as an estimate of downstream research performance. Future work should expand coverage to a larger range of specialized workflows and scientific domains. Ideally, benchmark performance could even be correlated with the results of deployment studies in live research settings. Such studies could establish a tighter relationship between the progression of model capabilities and whether AI systems actually improve scientific productivity.

## Research collaborators

Amelia Liu, Andrew Ho, Anne Marie Droste, David Martin, Edmund Wong, Edward Zhou, Isabelle Zhou, Joshua Park, Joy Jiao, Katie-Rose Skelly, Kenny Kim, Kevin Rao, Masatoshi Uehara, Max Marion, Nicole Fitzgerald, Rachel Dias, Suyash Shringarpure, Yuan Yuan, and Yunyun Wang.

## Expert scientist contributors

We thank the expert scientist contributors coordinated through Tacit Co. for authoring, reviewing, and validating LifeSciBench tasks and rubrics. Their contributions were essential to grounding the benchmark in realistic life science research work. Individual contributor names are not listed at the request of the vendor. Inclusion in this acknowledgement does not imply endorsement of the research, results, or conclusions.

## Appendix

### A Disclosures

#### A.1. AI Disclosure

We used AI tools to support literature review, language refinement, and routine engineering workflows during the development of this work.

#### A.2. Expert Contributor Disclosure

LifeSciBench tasks were authored, reviewed, and validated by external domain experts with relevant life-science training and experience. Expert contributors were compensated for their work.

#### A.3. Evaluation and Grading Disclosure

Model outputs were evaluated using task-specific rubrics developed during benchmark construction. Automated or model-assisted grading, where used, was applied against these rubrics rather than free-form preference judgments.

#### A.4. Institutional Disclosure

LifeSciBench was developed by OpenAI, and the evaluated systems include OpenAI models. Results should be interpreted with this institutional context in mind.

#### A.5. Data Availability and Safety Disclosure

Public release of tasks, rubrics, artifacts, or evaluation materials may be limited by licensing, privacy, proprietary information, or biological safety considerations. During benchmark construction and release review, we excluded or restricted content where broader dissemination could create biological safety risks.

### B Additional Benchmark Details

This appendix provides the partial-credit view cited in the Results, additional views of the benchmark’s attachment distribution and artifact-associated performance gaps, and the complete task-level model-comparison figure.

**Figure 8:**
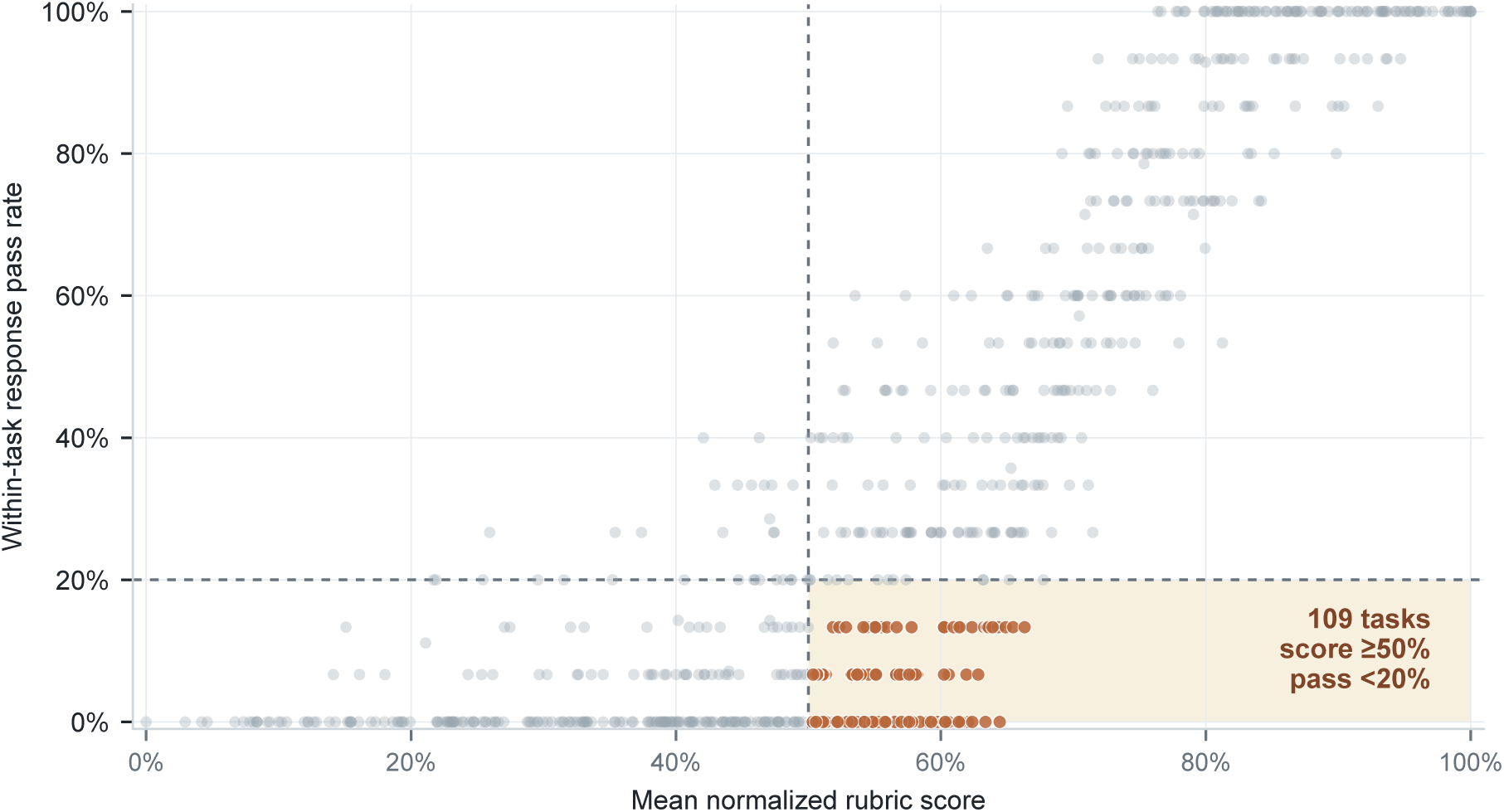
Partial credit versus task pass rate for GPT-Rosalind. Highlighted tasks have a mean normalized rubric score of at least 0.50 but a within-task response pass rate below 0.20, identifying substantial rubric credit without frequent passing responses.

#### B.1. Data Source & Evidence Distribution

**Figure 9:**
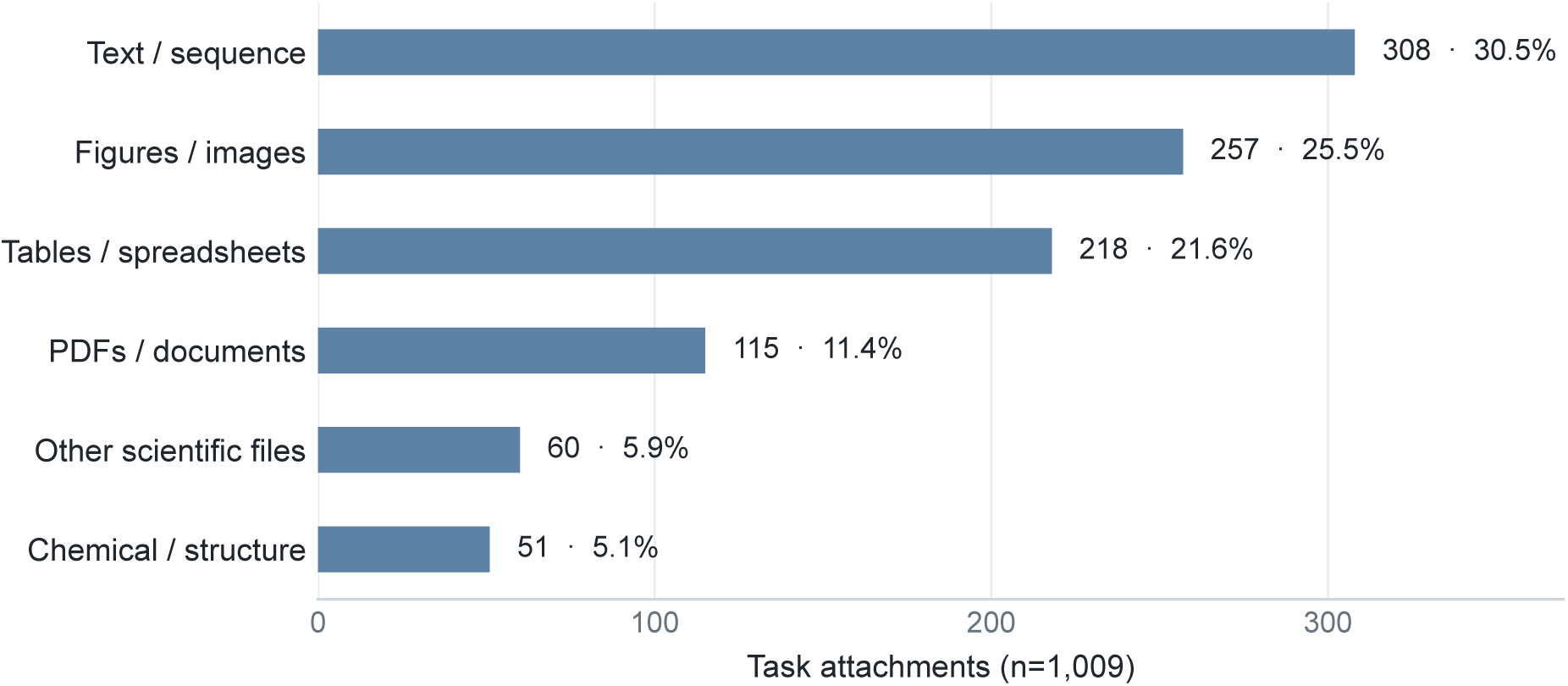
Distribution of task attachments by file category. The 1,009 attachments supplied with benchmark tasks are grouped by file category; labels report each category’s count and share.

#### B.2. Artifact-Associated Performance Gaps

**Figure 10:**
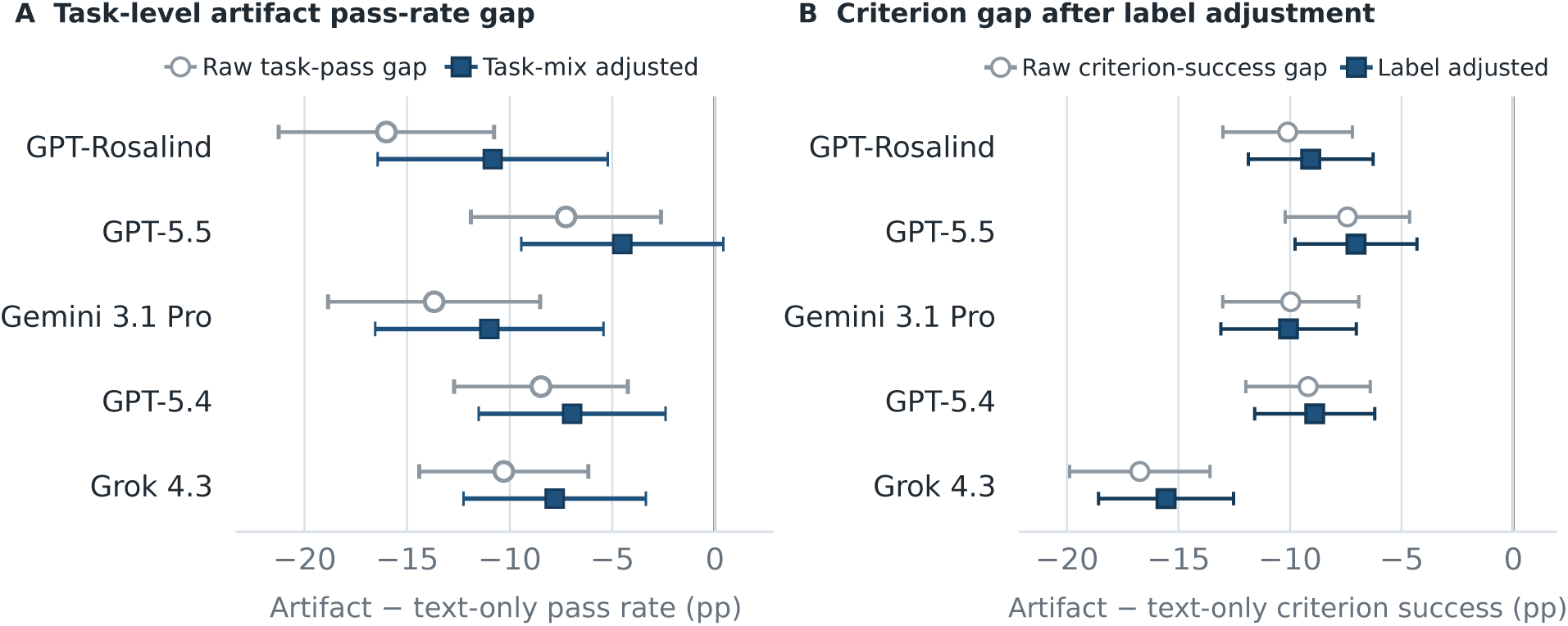
Artifact-associated performance gaps after adjustment for observed task composition. Adjusting for task type and criteria type yields a persistent performance gap between artifact-bearing tasks and text-only tasks. Models were fit separately for each evaluated system; circles show raw artifact-minus-text differences and squares show covariate-adjusted differences in units of percentage points. (A) Let pi denote the within-task mean response pass indicator and Ai = 1 denote a task with attached artifacts. The raw and adjusted task models were pi = α + βrawAi + εi and pi = α + βadjAi + γTXi + εi, where Xi contains workflow; the task-dominant operation, answer-form, exactness, and reasoning labels; rubric-size bin; and prompt-length quartile. Ordinary least-squares fits used HC3 robust standard errors. (B) Let sij denote mean criterion success for criterion j in task i. The raw and adjusted criterion models were sij = α + βrawAi + εij and sij = α + βadjAi + δTZij + εij, where Zij contains workflow and the criterion’s operation, answer-form, exactness, and reasoning labels. Weighted least squares gave each task equal total weight, with standard errors clustered by task. In both panels, points report 100βraw or 100βadj in percentage points and whiskers show 95% confidence intervals; negative values indicate lower performance on artifact tasks. Four models included all 750 tasks, whereas Grok 4.3 included 749.

#### B.3. Task-Level Model Comparisons

**Figure 11:**
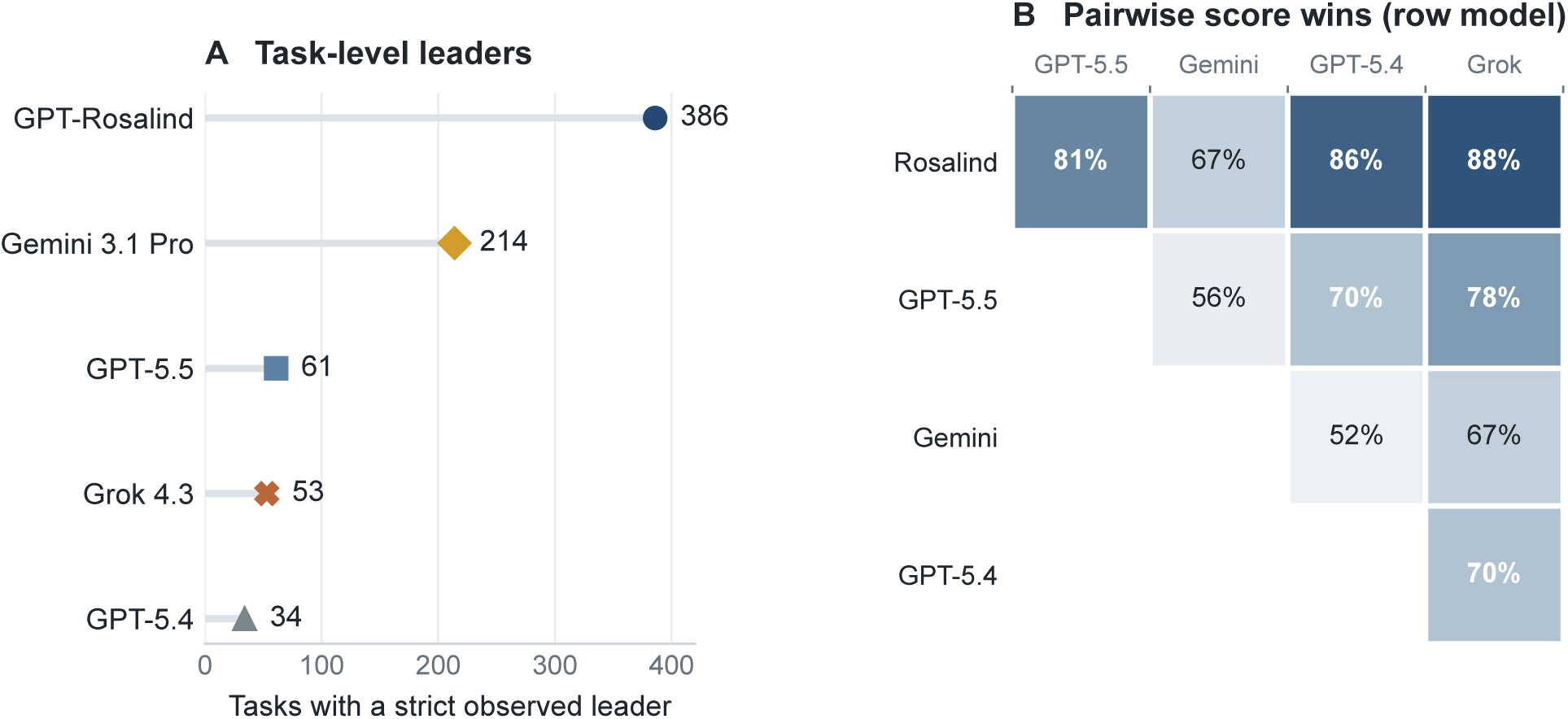
Task-level score comparisons across models. The left panel counts strict observed task-level score leaders; two tied tasks are excluded. The upper-triangular table-graphic shows each non-redundant model pair once and reports the fraction of non-tied task comparisons on which the row model scored higher than the column model. Comparisons involving Grok use 749 shared tasks before ties are excluded; all other pairs use 750. These descriptive comparisons use observed task-level mean scores and do not incorporate uncertainty in the within-task estimates.

#### B.4. Criterion-Level Rubric Taxonomy

Each of the 19,389 rubric criteria received one operation label and one answer-form label. Operation describes the type of action required by a criterion, whereas answer form describes the kind of output the response must provide. Counts can be found in Table 3. Task counts are distinct within each category and do not sum to 750 because a task commonly contains criteria from multiple categories.

**Table 3:** Criterion-level operation and answer-form taxonomy. Each rubric criterion receives one label on each axis. Criterion counts sum to 19,389 within each axis. Task counts are non-additive because individual tasks commonly contain criteria from multiple categories.

| Category | Plain-language description | Criteria | Tasks |
| --- | --- | --- | --- |
| **Operation: what the response must do** |  |  |  |
| Extract / identify | Recover or name a requested fact, entity, feature, or value. | 4,641 | 538 |
| Explain mechanism | Explain a causal, mechanistic, or conceptual relationship. | 2,887 | 535 |
| Critique / validate | Assess correctness, validity, quality, limitations, or evidentiary sufficiency. | 2,359 | 437 |
| Quantify / compute | Calculate or report a requested numerical quantity. | 1,784 | 290 |
| Interpret evidence | State what supplied evidence supports, contradicts, or implies. | 1,700 | 345 |
| Predict / infer | Infer an unobserved conclusion or predict an outcome. | 1,603 | 350 |
| Compare / rank | Compare alternatives, characterize a relation, or place items in order. | 1,289 | 337 |
| Recommend / decide | Select or recommend an action, option, or conclusion. | 958 | 284 |
| Generate / construct | Produce a specified scientific object, representation, or construct. | 922 | 125 |
| Design experiment | Specify an experiment, control, measurement, or discriminating test. | 881 | 199 |
| Enumerate / coverage | Enumerate a required set or cover multiple specified components. | 255 | 138 |
| Format / citation | Meet an explicit output-format or citation requirement. | 110 | 30 |
| **Answer form: what the response must return** |  |  |  |
| Explanation | Explanatory or justificatory prose. | 7,858 | 665 |
| Entity identifier | A named entity or other specific identifier. | 3,925 | 560 |
| Numeric | A number, range, statistic, or calculated quantity. | 1,801 | 277 |
| Sequence / structure | A biological sequence, chemical structure, or construct representation. | 1,332 | 145 |
| Plan / procedure | An experimental plan, protocol, action, or procedural step. | 1,322 | 300 |
| Boolean / category | A yes/no judgment or categorical label. | 1,268 | 285 |
| Relation / ranking | A comparison, ordering, or stated relation between entities. | 1,208 | 316 |
| List / set | A collection of multiple requested items. | 344 | 133 |
| Mixed | A response combining more than one answer form. | 331 | 136 |

### C Rubric Writing Guidelines

Rubrics were written according to the following principles:

1. **Specificity:** each criterion should describe a concrete property of the response.
2. **Atomicity:** each criterion should evaluate a single claim, calculation, decision, or constraint.
3. **Evaluability:** criteria should be answerable as satisfied or not satisfied from the model response alone.
4. **Grounding:** criteria should be supported by the task prompt, provided artifacts, accepted scientific facts, or expert consensus.
5. **Non-redundancy:** criteria should avoid double-counting the same requirement unless the repeated criterion captures a distinct aspect of the response.
6. **Operational usefulness:** rubrics should reward responses that are not merely correct in isolation but useful for the scientific decision posed by the task.

### D Example Tasks

#### D.1. Design, Optimization & Prediction - Golden Gate Cloning

**Figure 12:**
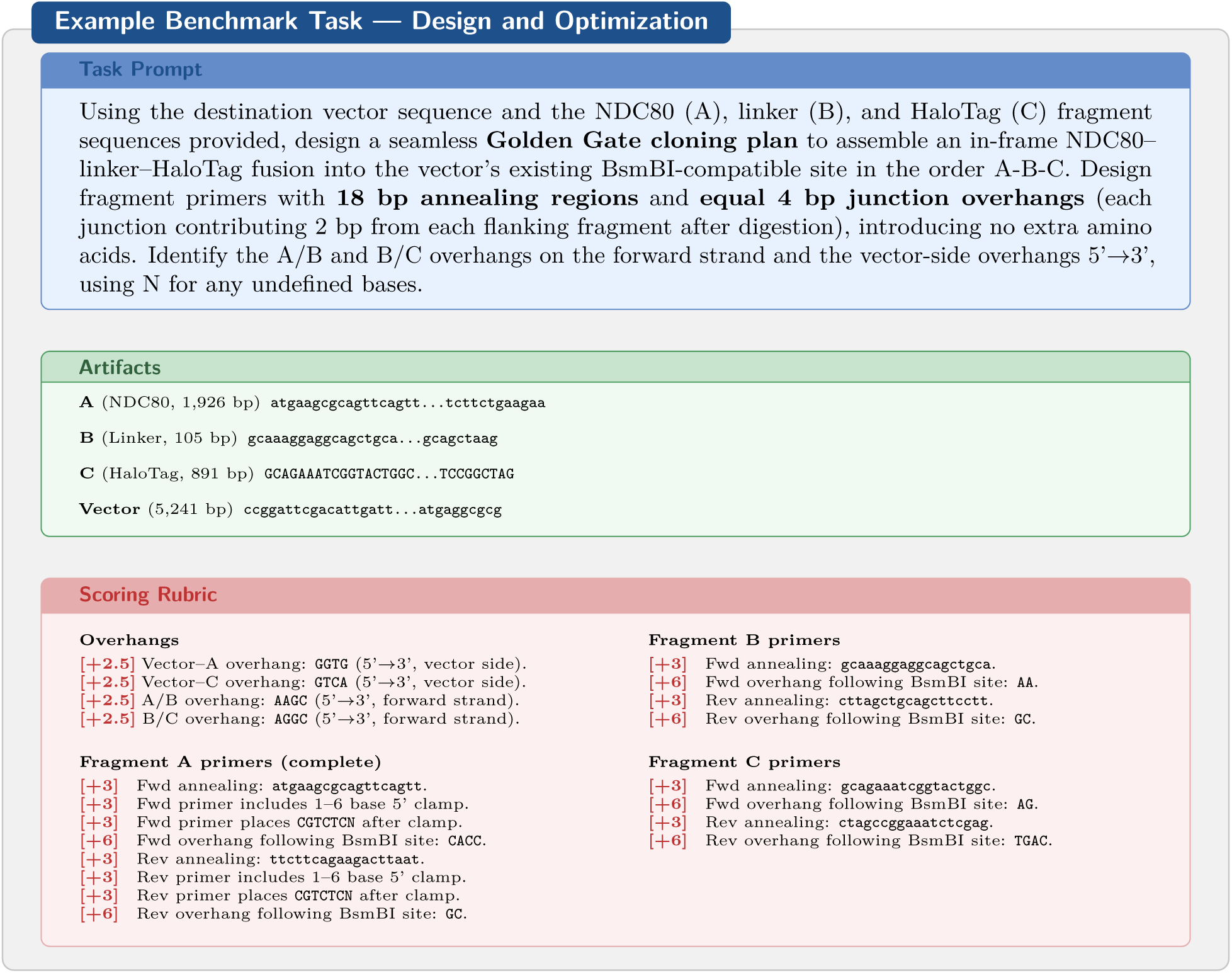
The model designs a three-fragment Golden Gate assembly using BsmBI, deriving junction overhangs from sequence context and producing complete primer architectures. Fragment sequences are truncated; full sequences are provided in the task. Fragment B and C rubric items show only annealing and overhang criteria; 5’ clamp and BsmBI site architecture items (+3 pts each) are omitted for brevity.

#### D.2. Evidence Handling - Plasmid Sequencing

**Figure 13:**
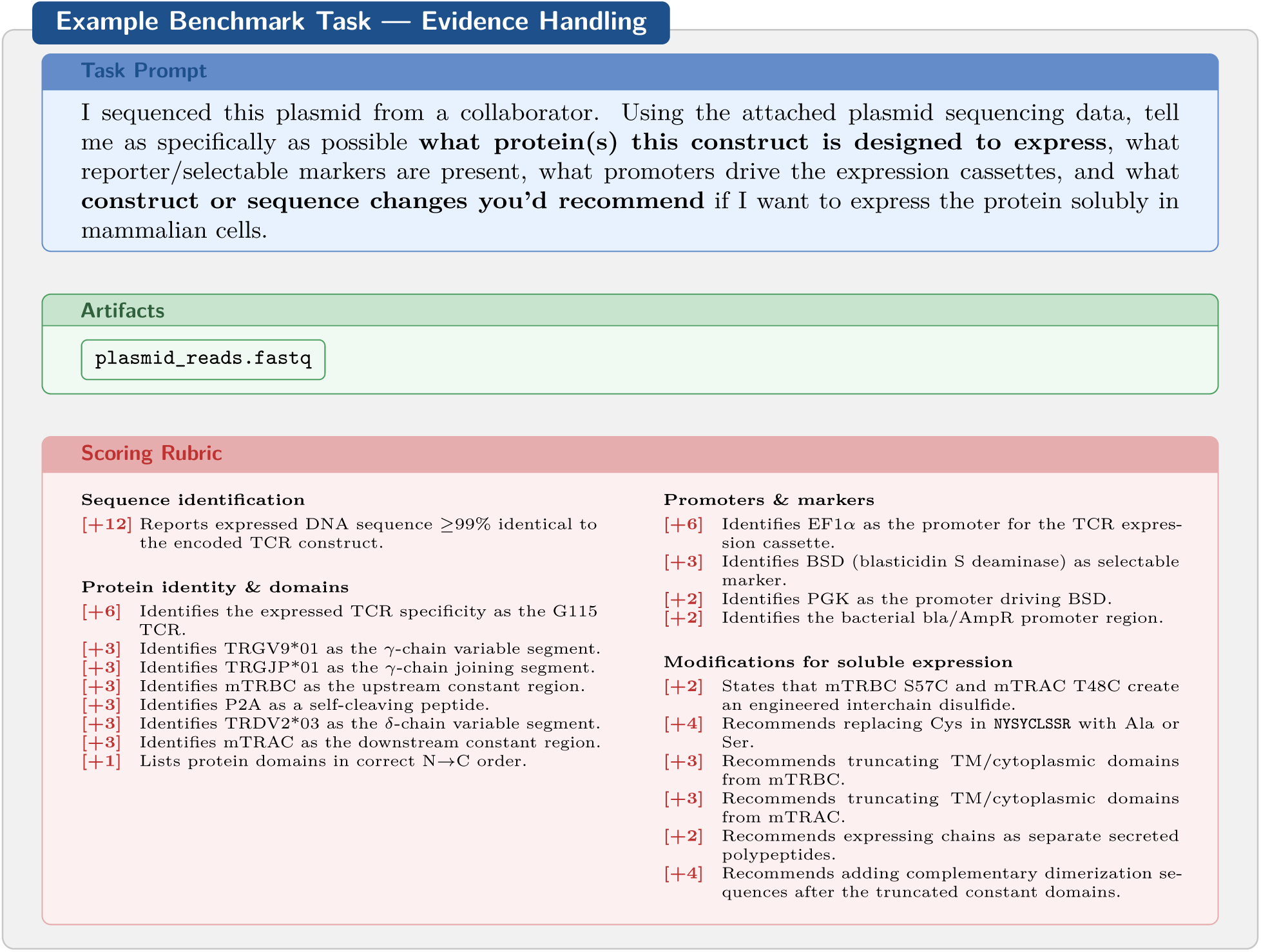
The model must assemble a complete construct description from raw sequencing reads, identifying TCR chain architecture, regulatory elements, and markers, then recommend specific molecular modifications for soluble expression. A representative subset of rubric criteria is shown.

#### D.3. Reasoning - Hippocampal Panel

**Figure 14:**
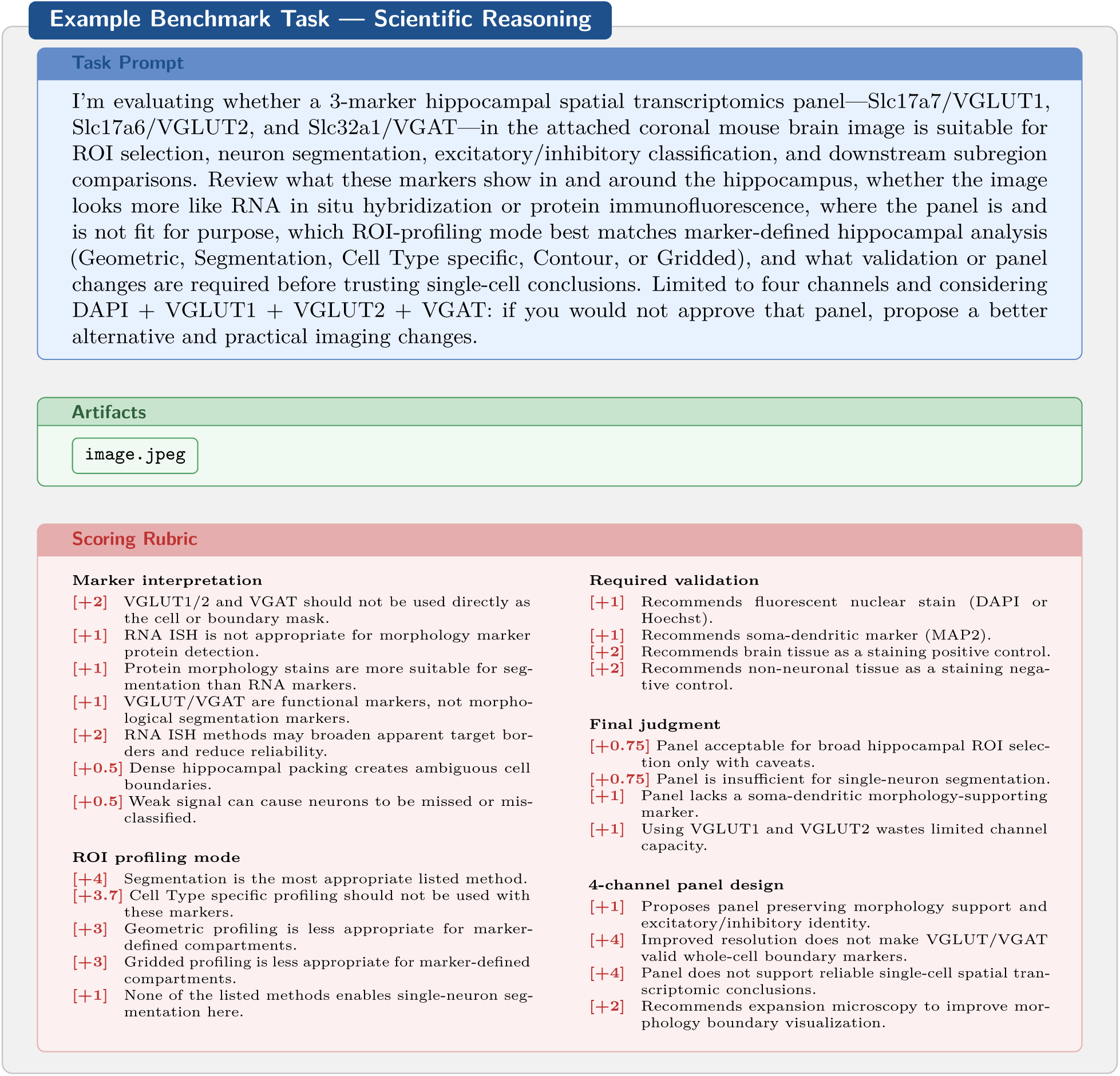
The model must integrate image evidence, marker biology, and profiling-mode constraints to assess a hippocampal spatial transcriptomics panel and propose panel improvements. The full rubric contains over 50 criteria spanning marker interpretation, ROI mode selection, required validation, final judgment, and 4-channel panel design; a representative subset is shown.

#### D.4. Translation - PILRA Xenograft

**Figure 15:**
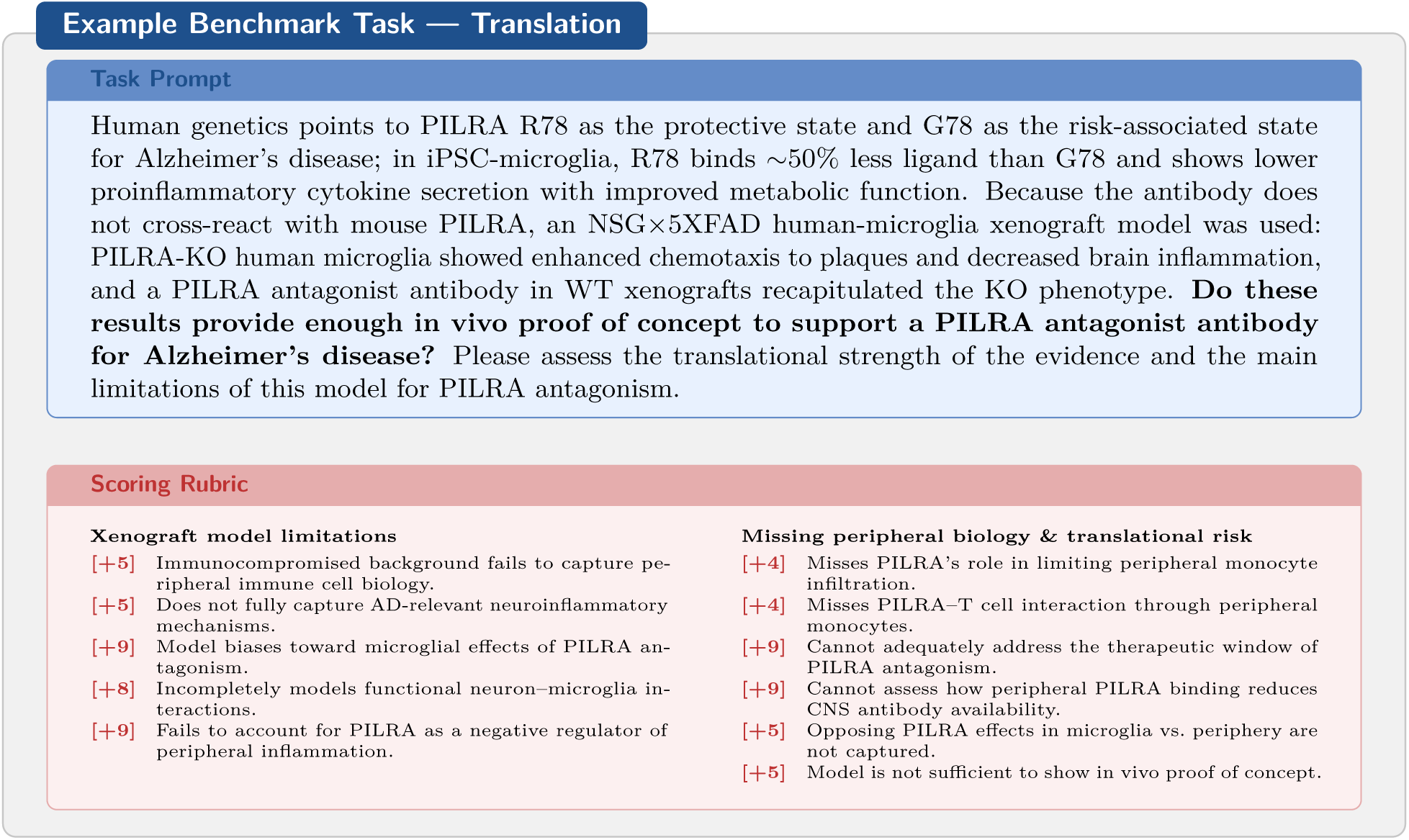
The model must critically assess the translational adequacy of a xenograft proof-of-concept study for a CNS-targeting antibody, identifying mechanistic gaps between the model system and the human disease context. A representative subset of rubric criteria is shown.

#### D.5. Validation & Operations - DNA Methylation NASH

**Figure 16:**
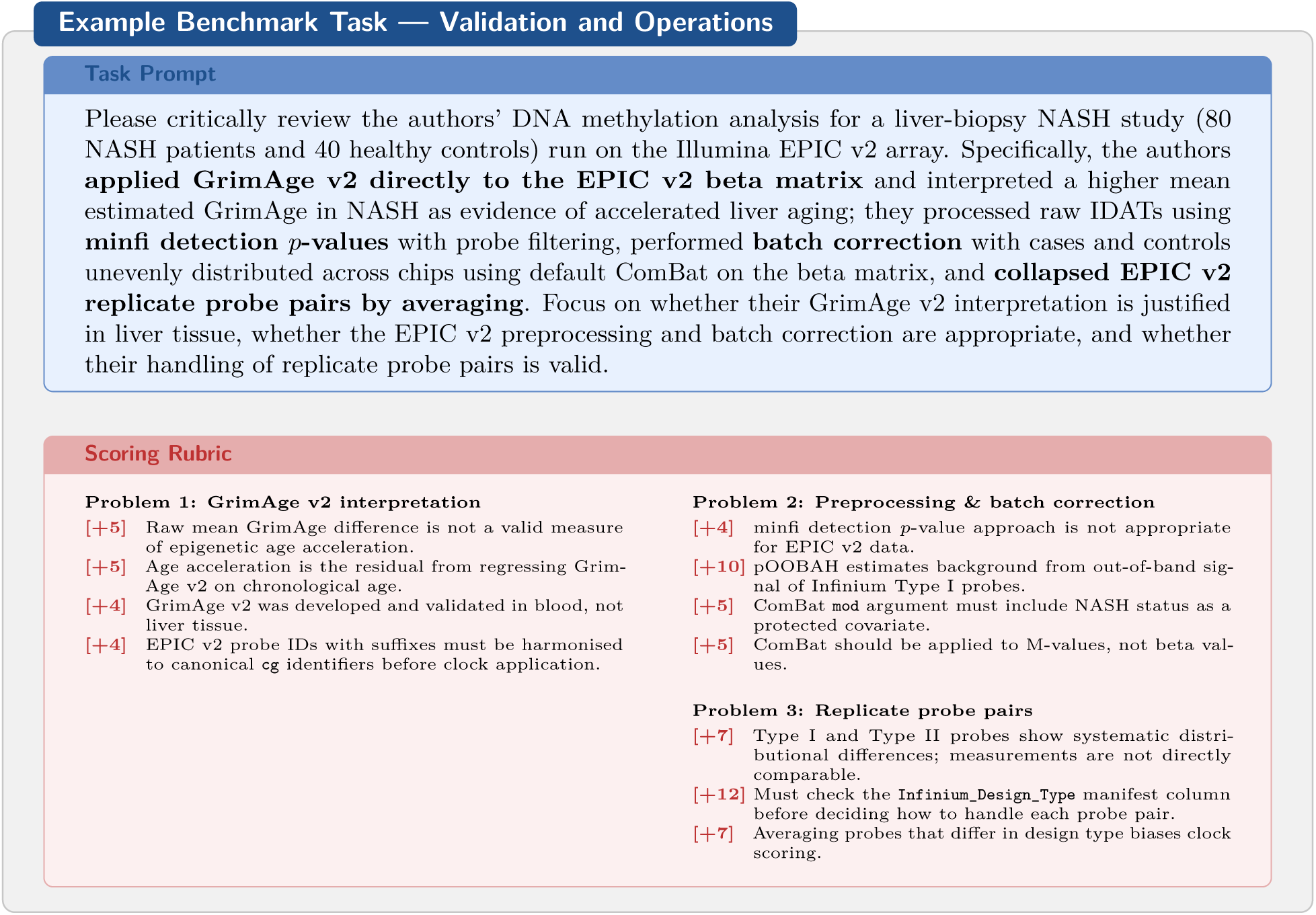
The model must identify three distinct methodological errors in a DNA methylation study: clock misapplication to non-blood tissue with incorrect acceleration calculation, inappropriate EPIC v2 probe quality control and batch correction, and invalid probe-pair averaging. A representative subset of rubric criteria is shown.

## Footnotes

1 Grok 4.3 had valid results for 749 of 750 tasks; its aggregate statistics are therefore computed over those 749 tasks.

## Notes

### Competing Interest Statement

LifeSciBench was developed by OpenAI, and the evaluated systems include OpenAI models. External expert contributors were compensated for their work.

